# Gcn5 and Ubp8 dependent ubiquitylation affects glycolysis in yeast

**DOI:** 10.1101/670885

**Authors:** Antonella De Palma, Giulia Fanelli, Elisabetta Cretella, Veronica De Luca, Valentina Panzeri, Valentina Roffia, Michele Saliola, Pier Luigi Mauri, Patrizia Filetici

**Author notes:** Corresponding Author: (PF). Department of Medical Surgical Sciences and Translational Medicine, Sapienza University of Rome, 00189 Rome, Italy (Valentina Panzeri).

## Abstract

Many of the molecular mechanisms affected by ubiquitylation are highly conserved from yeast to humans and are associated to a plethora of diseases including cancers. To elucidate the regulatory role of epigenetic factors such as the catalytic subunits of SAGA complex, KAT-Gcn5 and Ub-protease Ubp8, on ubiquitylation of non-histone proteins we have performed a comprehensive analysis of the Ub-proteome in yeast *Saccharomyces cerevisiae* in strains disrupted in Gcn5, Ubp8 or both respect to wild type. We found significative alteration of ubiquitylation in proteins belonging to different functional categories with a recurrence of identical proteins in absence of Gcn5 or Ubp8 indicating shared targets and their interlaced function. Among the processes involved we noteworthy identified all major enzymes engaged in energy metabolism and glycolysis such as PFK1, PFK2 and others showing increased ubiquitylation respect to WT. We showed that the higher degree of ubiquitylation found is at post-translational level and does not depend on transcription. Noteworthy, we found *in vivo* severe defects of growth in poor sugar medium and inability to adaptive switch from fermentative to respiratory growth in strains lacking Gcn5 and Ubp8. Our findings data provide a novel, direct link, between metabolism and epigenetic control with a novel role of DUB-Ubp8 and KAT-Gcn5 on the ubiquitylation marking all the main glycolytic enzymes required for an effective execution of the glycolytic flux. Collectively our experimental results and the proposed model can lead to future research and innovative strategies that by targeting epigenetic modulators might be able to lower sugar utilization also in human cells.

**Author Summary:** Molecular mechanisms dissected in simple yeast might be translated to similar circuitries in human cells for new discoveries in human diseases including cancer. Ubiquitylation of proteins is an evolutionary conserved mechanism required for many biological processes. Different post-translational modifications (PTMs) such as ubiquitylation, acetylation, methylation etc. are reciprocally regulated for deposition or removal. Epigenetic factors writing the PTMs code are often components of multiproteic complexes such as SAGA complex that holds the K-acetyltransferase (KAT) Gcn5 and the Ubiquitin-protease (DUB) Ubp8 highly conserved in Evolution. Cells respond to environment and nutrients by changing metabolism and group of enzymes involved in specific pathways are often coregulated by the deposition of selected PTMs. This study analyses the composition and quantitation of Ub-proteins differentially modified in absence of KAT-Gcn5 and DUB-Ubp8 in yeast. Interstingly, we highlighted the role of Gcn5/Ubp8 dependent ubiquitylation in marking major glycolytic enzymes necessary for glucose utilization. Our study suggests a novel regulatory pathway and, considering that lowering glycolysis is a promising strategy to target tumor metabolism, we propose this study as an interesting perspective to lower enhanced glycolysis in tumors.

## Introduction

Conservation in Evolution allows translation of discoveries obtained in simple models useful for the interpretation of more complex issues in human cells. Accordingly, the simple budding yeast is one of the best models to study molecular mechanisms regulating complex networks and circuitries [1]. Proteins are differentially regulated by a plethora of post-translational modifications (PTMs) often involved in a reciprocal cross talk and regulation. Lysine acetylation, for example, prevents further poly-ubiquitylation on the same residue with a direct impact on several functions including protein turn over and proteasomal degradation [2]. The big multiproteic SAGA complex (Spt-Ada-Gcn5-acetyltransferase) is an epigenetic key regulator of acetylation and establishes the transcriptional competence of gene promoters through the opening of chromatin to the transcriptional machinery [3]. Its relevance is not merely confined to histones modifications but extend far behind to the deposition of PTM marks on non-histone proteins. It is composed of 21 widely conserved proteins grouped into functional submodules, it harbours K-acetylatransferase KAT2-Gcn5 mediating acetylation and Deubiquitylase (DUB) Ub-protease Ubp8 mediating the deubiquitylation [4, 5]. The coexistence in SAGA complex of acetylation and deubiquitylation modules sustains their possible interdependency and interlaced functions. Gcn5 and Ubp8 indeed showed functional interaction in fermentative or respiratory growth conditions [6] and at high and low temperatures previously reported to be correlated with glycolysis [7]. In the present study we investigated polyubiquitylated proteins expressed in absence of KAT-Gcn5, DUB-Ubp8 or both in comparison with wild type strains. We approached this study by expressing His6-Ub in the different yeast strains performing selective biochemical purification of tagged Ub-proteins. After elution we analyzed the composition of Ub-proteins obtained with a gel-free and label-free proteomic approach based on the coupling of micro-liquid chromatography and tandem mass spectrometry (µLC-MS/MS). Certainly, proteomics is proving its potential as tool of choice to reach a deepened characterization of protein-protein interactions and PTMs that regulate crucial biological processes. Using a multidimensional chromatographic system coupled to mass spectrometry it was investigated, for example, the function of TFIID in *S.cerevisiae* by identifying new connections between its components and novel subunits of SAGA complex obtaining the most complete proteomic interaction map of an eukaryotic transcription machinery [8]. Recently, Wilson M.A. et al. [9] demonstrated with a proteomic approach that SAGA complex also deubiquitylates SNF1 altering its kinase activity confirming that the acetylation of non-histone proteins plays a direct role in the regulation of protein functions. This evidence is in agreement with results obtained in a study on Gcn5 and Esa1 mutants [10] with SILAC and found their coordinated function in the regulation of cell growth and division. It was reported that, PTMs regulate central carbon metabolism, glycolysis and fermentation where a high number of key enzymes engaged in sugar utilization and energy supply were marked by phosphorylation, acetylation and ubiquitylation indicating a direct involvement of PTMs in the regulation of their catalytic activity [11]. In addition, we have recently demonstrated that Gcn5 and Ubp8 beside their role as catalytic subunits of SAGA complex, are also involved in the control of the respiratory metabolism and mitochondrial functions [6, 12]. Following this line of reasoning and starting from a shotgun proteomic profiles and *in vivo* phenotypic assays in strains missing Gcn5 and Ubp8 or both we wanted to deeply characterize the composition of differentially ubiquitylated proteins in order to clarify their role and investigate possible correlation with the regulation of sugar metabolism. Our experimental data provided significative results thus demonstrating an altered ubiquitylation on main glycolytic enzymes in absence of Gcn5 and Ubp8. We believe that our results shed light on a novel role of SAGA complex in the ubiquitylation of metabolic enzymes opening to new perspectives on the relevance of ubiquitin in regulating glucose metabolism also in higher cells. Therefore, the modulation of PTMs on key enzymes may lead to a new strategy to slow or block sugar metabolism that may counteract the rapid and energy demanding proliferation of cancer cells.

## Results

### Genetic and functional interaction between HAT-Gcn5 and DUB-Ubp8 in *S.cerevisiae*

Based on the coexistence in the multiproteic SAGA complex of the KAT and DUB modules and the deubiquitylation activity of Ubp8 on the centromeric histone variant Cse4 previously reported [12] we wanted to analyse whether the global process of protein ubiquitylation might be dependent on Gcn5 or Ubp8. In *S.cerevisiae*, the functional interaction between genes is tested at genetic level by comparison of the growth of single disrupted strains respect to the strain carrying the double genes disruption in order to analyze the resulting synthetic phenotype and the fitness of the strains grown in selective conditions (Fig. 1A). We previously demonstrated the genetic interaction between Gcn5 and Ubp8 in respiratory growth conditions [6, 12]. Here we have assayed *S.cerevisiae gcn5Δ, ubp8Δ* and *ubp8Δ/gcn5Δ* strains and confirmed their genetic interaction by growing at low (12°C), physiological (28°C) and high (35°C) temperatures (Fig. 1B). Growth spot assay suggested that there is a severe growth defect at 12°C and at 35°C with a remarkable synthetic defective phenotype in the growth of the double disrupted *ubp8Δ/gcn5Δ* strain. Based on these genetic evidences we decided to investigate if loss of Ubp8 and Gcn5 or both might, possibly, affect the ubiquitylation process of non-histone proteins in an interdependent way. *Purification of polyubiquitylated proteins in WT, ubp8Δ, gcn5Δ and ubp8Δ/gcn5Δ strains.* The 6His-Ub-proteins were expressed in WT, *ubp8Δ, gcn5Δ and ubp8Δ/gcn5Δ* strains according to procedure described in material and methods (Fig. 2A), purified on Ni-column [13] and analyzed on western blot. Fig. 2B shows the selective purification of the obtained His6Ub-proteins, evidenced with anti-His6 antibody (Fig. 2B). Unpurified, total protein extracts, indicated as input, were hybridized with anti-Ada2 antibody as an internal control indicating the expression of a constitutively expressed protein in our cellular extracts. The quality of the Ub-protein obtained prompted us to go further and perform a proteomic characterization after trypsin digestion of 6His-Ub-proteins retained by means of a shotgun label-free platform. In particular, our proteomic approach, based on a µLC-MS/MS, allowed the identification of 447 distinct proteins, whose distribution in the four analyzed strains is represented by a Venn diagram shown in Fig. 2C. A total of 16 µLC-MS/MS runs were acquired, evaluating two preparations for the four experimental conditions (biological replicates) and analyzing them in duplicate (technical replicates). The complete list of proteins identified for each group is reported in Table S1. Considering the strains disrupted in KAT-Gcn5, in the Ub-protease Ubp8 or in both (*gcn5 Δ ubp8 Δ and ubp8 Δ/gcn5 Δ*) compared to WT, we observed that the majority of proteins identified in *ubp8Δ* and *gcn5Δ* strains were also found in the double *ubp8Δ/gcn5Δ* strain. In fact 193 proteins, corresponding to the 43,2% of total, were shared among the four strains, suggesting that these proteins represent common targets of both, Gcn5 and Ubp8. Moreover, 48 proteins have been identified in common between WT and *ubp8 Δ* and 18 between WT and *gcn5 Δ*, suggesting that Ubp8 is involved in the regulation of more proteins respect to Gcn5 (10,7% vs 4%). This observation was expected and based on the Ub-protease activity of Ubp8, whereas we expect that Gcn5 acts indirectly possibly through the deposition of a counteracting acetyl group on specific lysines. Based on the general results obtained and recapitulated in the Venn diagram (Fig. 2C), we decided to investigate in more depth the differences in expression of ubiquitylated proteins in the examined strains by performing label-free differential analyses.

**Fig. 1.**
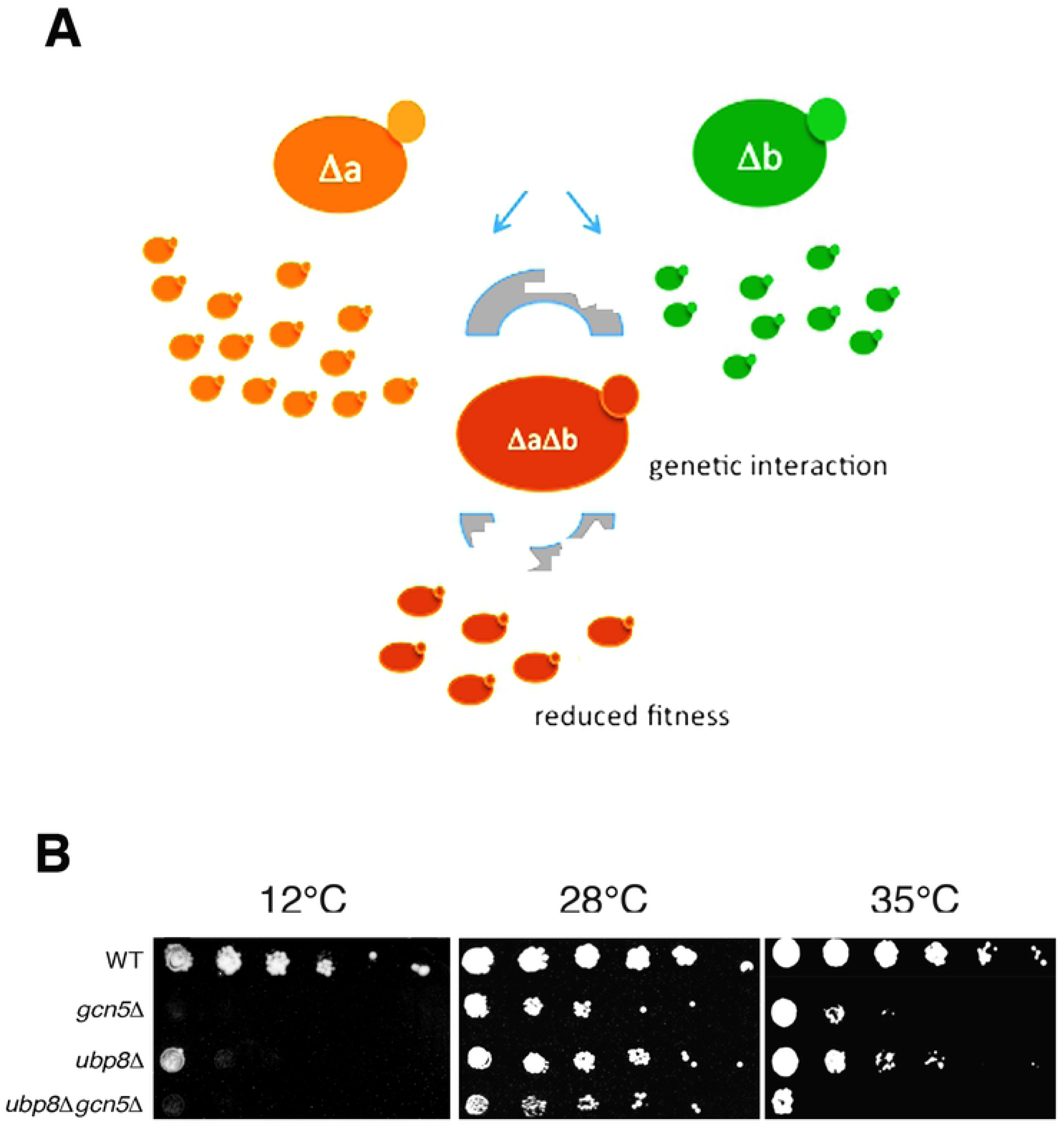
Gcn5 and Ubp8 show genetic interaction. A, schematic diagram showing the effects on a strain fitness in case of genetic interaction between two genes. B, Growth spot assay of *ubp8Δ, gcn5Δ and ubp8-gcn5Δ* strains in comparison with WT grown on rich medium in cold (12°C) physiological (28°C) and high temperature (35°C).

**Fig. 2.**
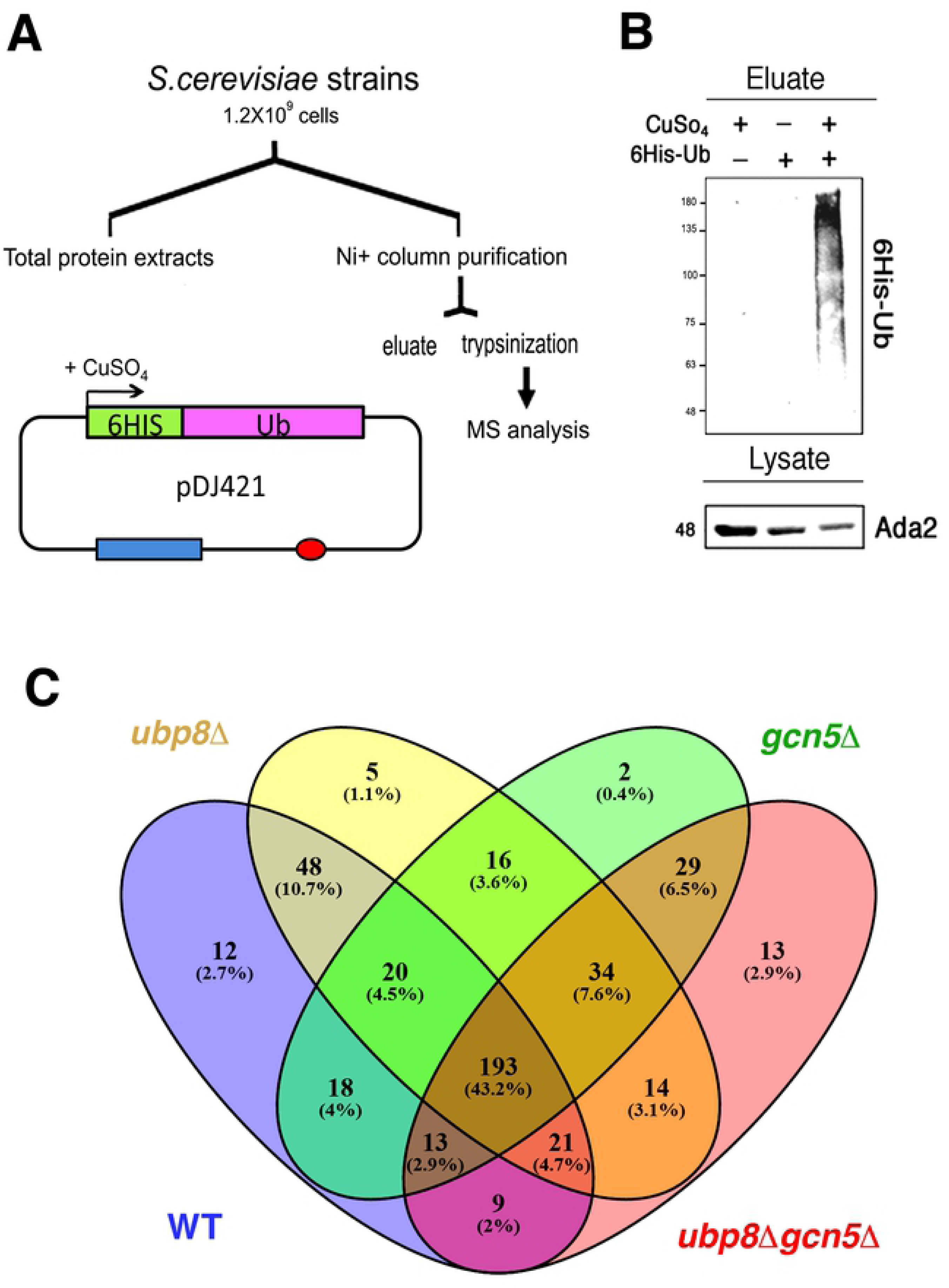
Expression and purification of His6-Ub proteins in *S.cerevisiae*. A, Schematic protocol for the expression of His6Ub proteins in strains containing pDJ421. His-Ub were expressed in CuSO^4^, purified through Ni^+^-column and analyzed by MS after trypsinization. B, Western blot analysis showing the eluate of His6-Ub proteins respect to the controls hybridized with anti-His6 antibody, total lysates were probed with anti Ada2 antibody as internal standard. C, Venn diagram of proteins distribution found in WT (blue), *ubp8Δ* (yellow) *gcn5Δ* (green) *and ubp8-gcn5Δ* (red). Area of intersection contain proteins common to different strains.

### Label-free differential analysis

Using MAProMa software [14] all the sixteen protein lists were aligned and for each strain a unique list has been created averaging the spectral count values (SpC*) of the identified proteins in order to estimate their relative abundance in the pairwise comparisons with *gcn5 Δ, ubp8 Δ* and *ubp8 Δ*/*gcn5 Δ* strains respect to the WT strains. To this end, the two algorithms of MAProMa software, DAve and DCI, were applied, representing the ratio and the confidence in differential expression, respectively, for each protein between two samples. Using stringent filters described in M&M section (see pag. XX) to maximize the confidence of identification, a total of 107 proteins were found differentially expressed and reported in Table 1 with selected details and in Suppl. Table S1 in extended form. In particular, 6 and 46 proteins resulted respectively down- and up-represented in *ubp8 Δ* compared to WT, while, 48 proteins were more abundant in *gcn5 Δ* compared to WT and 8 were down-represented in *gcn5 Δ* strain. Finally, 12 and 64 proteins resulted down- and up-represented, respectively in *ubp8 Δ*/*gcn5 Δ* strain compared to WT. In addition, known genetic (G) or physical (P) interactions with, respectively, Ubp8 or Gcn5 are reported, and, finally, the corresponding name of the human hortholog gene. The trend of the abundance of proteins among the pairwise comparisons of the three mutant strains respect to WT is shown through a color code corresponding to the individual DAve values assigned: for DAve values ranging from +0.40 to +2.00 color gradations are used from light red to dark red and for those ranging from −2.00 to −0.40 chromatic shades from dark blue to light blue. Moreover, in order to give a more comprehensive view of the pathways and of the proteins involved with KAT-Gcn5 and DUB-Ubp8 interactions, in Table 1 a color code is assigned also to those proteins whose DAve and DCI values do not pass the filters in all the considered comparisons. So, proteins with DAve values ranging from −0.4 to +0.4 appear with light red and light blue coloring respectively, while the unidentified or unchanged ones are shown with white code. Therefore, observing the table and the trends of the differentially expressed proteins among the three comparisons, it is immediately evident that some proteins seems to be more ubiquitylated in absence of DUB-Ubp8 others of KAT-Gcn5 indicating the major regulatory role of Ubp8 and Gcn5 on the specific protein. For example, ASN2, LEU1, PYC2, TDH2, GSY2, PGM2 and TY1B-PR1 are proteins that equally change in the *gcn5Δ* and in the double mutant strain, these evidences suggest that they are closely related to Gcn5 as much more influenced by the mutation of this gene. In the same way, YEF3, CDC19, CYT1, HSC82 and YHR097C equally change in the *ubp8Δ* and in the double mutant strain, thus demonstrating that they are closely related to Ubp8 as much more influenced by the mutation of this gene. Another interesting aspect concerns those proteins (such as, EFT1, PGI1, GPD1, SOD1, AMD1 and URA2) that show a marked increase or reduction in their levels if we consider the *ubp8Δ*/*gcn5Δ* strain compared to the single ones, demonstrating in this case the synergistic effect exerted by the double deletion of Ubp8 and Gcn5. Finally, in Table 1 we can also observe other proteins (ILV6, PFK1, PFK2, SCH9, HNT1, GCV2) that show the same differential trend both in the single and in the double mutant strains. Noteworthy the physical (P) or genetic (G) interactions of Ub-proteins with either Gcn5 and Ubp8 supports these results, indicating a direct effect of Gcn5 and Ubp8 in their modification. Among these, we found proteins that can be grouped on the basis of their functional role, such as amino acids biosynthesis, glycolysis, fermentation, oxidative phosphorylation and energy metabolism. With the aim to better visualize the pathways and the biological relevant interactions in which these proteins are involved in, the STRING database [15] was queried to build a network of both known and predicted protein-protein interactions. Differentially levels of proteins were grouped in ten sub networks based on their molecular function and represented by nodes and grey edges indicating protein-protein interactions. Applying the same chromatic scale of Table 1, Figure 3 show the three networks involving the differentially expressed protein behaviors towards the pairwise comparisons of *gcn5Δ, ubp8Δ* and *ubp8Δ*/*gcn5Δ* strains against WT condition.

**Table 1.**
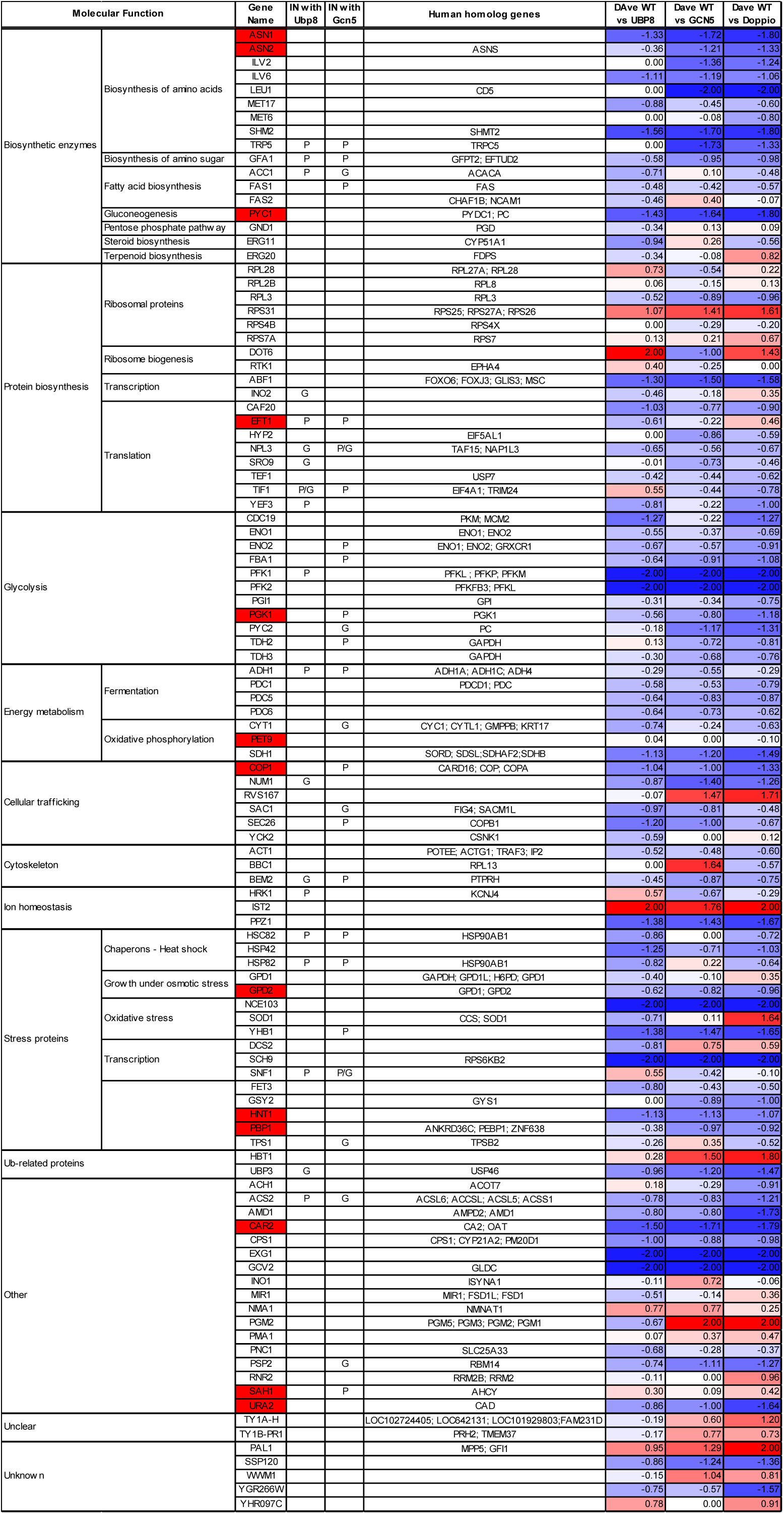
Ub-proteome of the three S.cerevisiae mutant strains, as determined by proteomic analysis.

**Fig. 3.**
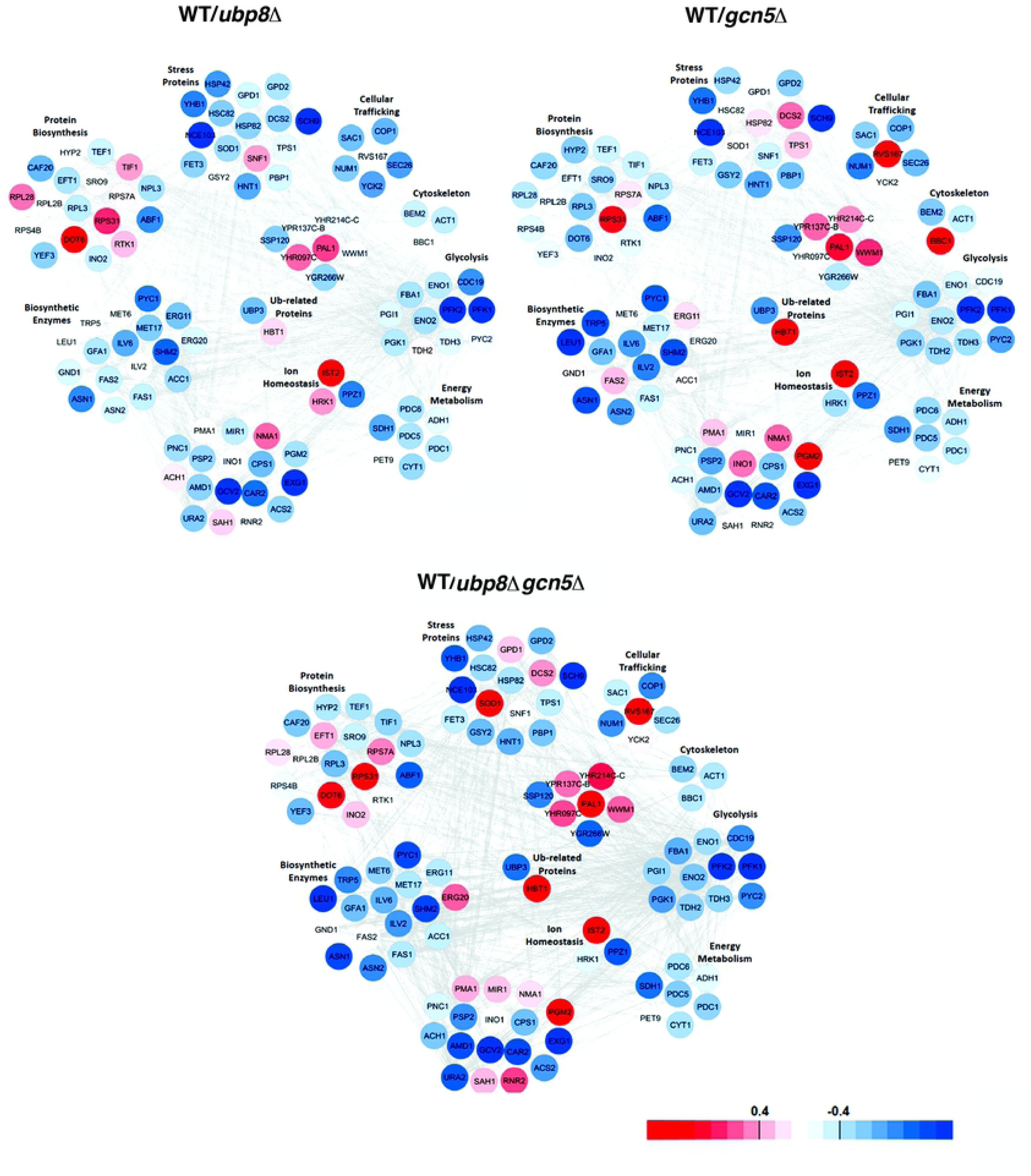
Interactome networks built using STRING database through the mapping of differentially expressed proteins identified comparing *ubp8*Δ, *gcn5*Δ and *ubp8*Δ-*gcn5*Δ strains versus WT condition. Each protein is identified by its Gene name (see Supplementary Table S2 for the complete reference), represented as a node and grouped to others belonging to the same pathway. Only experimentally and computationally predicted protein-protein interactions are considered and indicated as grey edges. The color code of distinct nodes represents the DAve value and the relevant chromatic scale (reported in figure) ranges from −2.00 to 0 (dark blue to white) and from 0 to +2.00 (white to dark red). Proteins with DAve ≥ l0.4l and a DCI ≥ l6l pass the filters and could be considered differentially expressed in the considered comparison.

Differentially expressed proteins resulted from the MAProMa comparison of *ubp8*Δ, *gcn5*Δ and *ubp8*Δ-*gcn5*Δ strains versus WT condition. In particular, each protein (identified in figure through its Gene Name) is marked by a color code which it is defined by the DAve value obtained in the three examined comparisons. The color is assigned according to a chromatic scale (reported in figure) which represents the confidence ranges of DAve values adopted (from −2.00 to 0 a gradient from blue to white and from +2.00 to 0 a gradient from red to white). Positive DAve values indicate proteins down-represented in mutant strains, while negative DAve values indicate proteins up-represented in mutant strains. Proteins are indicated for Molecular Function, Gene name, known physical (P) and/or genetic (G) interactions with Ubp8 and/or Gcn5 and the name of the human hortholog. In red the yeast genes corresponding to human horthologs related to human pathologies. The complete list of the reported proteins was extracted from the differential lists in Table S2.

### Major glycolytic enzymes are differentially ubiquitylated in absence of Gcn5 or Ubp8

The fact that several altered proteins found in our screening are linked to glycolysis, oxidative stress and energy metabolism (see Fig.3) could support the hypothesis that glycolysis is impaired in cells lacking Gcn5 and Ubp8. This observation, previously reported by Tripodi et al. [11], would strengthen our proteomic data and sustain a regulatory role of ubiquitylation on fundamental glycolytic enzymes. We report in Fig. 4A the scheme of the major steps composing the glycolytic flux for the utilization of glucose bringing to production of pyruvate from glucose-6-P. Major enzymes involved in these metabolic reactions were identified in our screening and Fig. 4A shows the proteins colored according to the color code previously described. This finding sustain a strong effect on the ubiquitylation level on the most important enzymes involved in glycolysis suggesting an impairment in the glycolytic flux progression. In Fig. 4B is reported the detailed color code of each identified enzyme found in strains disrupted in Gcn5, Ubp8 or both compared with the wild type. The panel provides the overall picture demonstrating how strong is the differential level of ubiquitylation on these proteins. We therefore asked whether the transcriptional expression was also affected by Gcn5 or Ubp8 disruption, and we made the analysis of the PFK1 and PFK2 mRNA expression, heavily ubiquitylated in all the strains and PYC1 showing only a slight difference between strains disrupted in Gcn5 or Ubp8. Fig. 4C shows diagrammed results obtained with RT qPCR on mRNA expression demonstrating no effects of either Gcn5 or Ubp8 disruptions on mRNA expression of the analyzed genes. Collectively, these results exclude any up-regulation at transcriptional level of these genes confirming that the higher degree of ubiquitylation found is at post-translational level and does not depend on transcription.

**Fig. 4.**
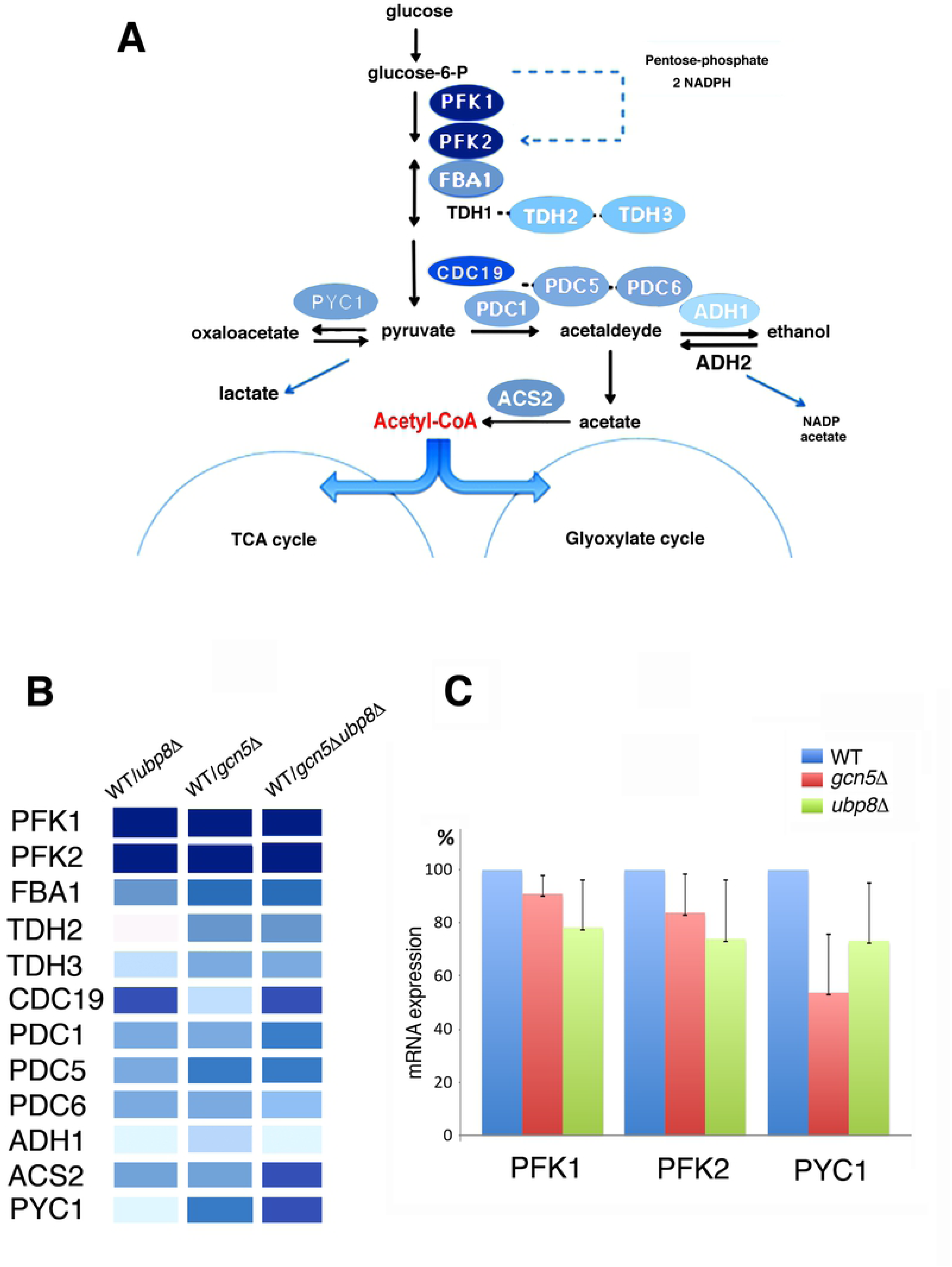
Major glycolytic enzymes are differentially ubiquitylated in absence of Gcn5 and Ubp8. A, Glycolytic pathway in *S.cerevisiae*, major glycolytic enzymes are shown according to the color code scale indicated in Fig.3. B, Detailed color code scale found in *ubp8Δ, gcn5Δ and ubp8Δ-gcn5Δ* compared with WT values for the glycolytic enzymes identified in our MS analysis. C, qPCR of PFK1, PFK2 and PYC1 mRNA expression in WT, *ubp8Δ* and *gcn5Δ* strains, respect to constitutive actin mRNA.

### Glycolysis is impaired in absence of Gcn5 and Ubp8

In order to assay their efficiency in sugar utilization, we then decided to grow WT, *gcn5Δ, ubp8Δ* and the double *ubp8Δ*/*gcn5Δ* strains in medium with high (2%) and low (0,1%) glucose in order to assay their efficiency in sugar utilization. According to our model, Fig. 5A shows a severe defect of growth in absence of Gcn5 and Ubp8 in low glucose that is not recovered by addition of downstream products such as lactate and ethanol. We repeated this analysis following prolonged growth in rich medium liquid cultures in the presence of 2% and 0,2% glucose. A strong impairment of growth in absence of Gcn5 and Ubp8 was observed after 24 hours in low sugar, while in the wild type there was only minor effects of sugar concentration on growth. Disrupted strains were unable to duplicate after 24 hours and growth was blocked, indicating inability to switch to respiratory conditions (Fig. 5B). To confirm these data, we used the Alcohol-dehydrogenase activities, Adh1 and Adh2, as respiro-fermentative glycolytic flux markers. In fact, Adh1, which is primarily involved in ethanol production, is expressed in fermentative, high glucose, conditions [16]. On the other side, Adh2 is activated when glucose is exhausted and the cell switches to respiratory metabolism by growing mainly on the accumulated ethanol. The transition from one condition to the other can be followed by the native in-gel appearance of the glucose-repressible ADH2 gene product with an Adh-specific assay, previously used to dissect the ADH genetic system [17] [18] [19]. To this end, protein extracts from 2% and 0,2% glucose-containing cultures of WT, *ubp8Δ, gcn5Δ* and *ubp8Δ*/*gcn5Δ* strains grown for 30, 60 and 90 hours were fractioned on native PAGE and stained for Adh. It must be underlined that the use of 0,2% glucose was a necessary condition to avoid the nearly complete impairment of growth of *gcn5Δ* and *ubp8Δ* strains in more stringent 0,1% glucose-containing medium. The Adh pattern of the wild type on 2% and 0,2% glucose can easily describe the expected growth phases (Fig. 5C). The exclusive presence of Adh1 in exponential phase indicated fermentative growth. Then, the progressive appearance of Adh2 in 2% glucose at 60 and 90 hours of growth indicated exhaustion of sugar and switch to respiratory conditions supported by the oxidation of the accumulated ethanol (Fig. 5C, lanes 1-3). The Adh pattern of the wild type in low glucose indicated a slight fermentative growth in log phase quickly shifted to respiration by the strong activation of Adh2 at 60 hours and highly reduced levels of both Adh versions at 90 hours caused by the exhaustion of all carbon sources (Fig. 5C, lanes 12-15). In agreement with the growth (Fig. 5A), the absence of Adh2 in both *gcn5Δ* and *ubp8Δ* strains confirmed their inability to grow in 0,2% glucose indicating a severe impairment in the activation of respiratory metabolism. Interestingly, the very small amount of Adh2 present in *gcn5Δ* culture in 2% glucose in all phases of growth, evidenced by the outlined scheme on the left of Fig. 5C, suggested the putative activation of a respiro-fermentative metabolism (Fig. 5C, lanes 4-6) [20]. On the contrary, this was not found in *ubp8Δ* cultures that displayed only Adh1 bands thus indicating no activation of respiratory metabolism. Although few differences between *gcn5Δ* and *ubp8Δ* strains, collectively these experimental data demonstrate that, in absence of Gcn5 and/or Ubp8, cells are unable to adapt their metabolism during the progression from fermentative to respiratory conditions.

**Fig. 5.**
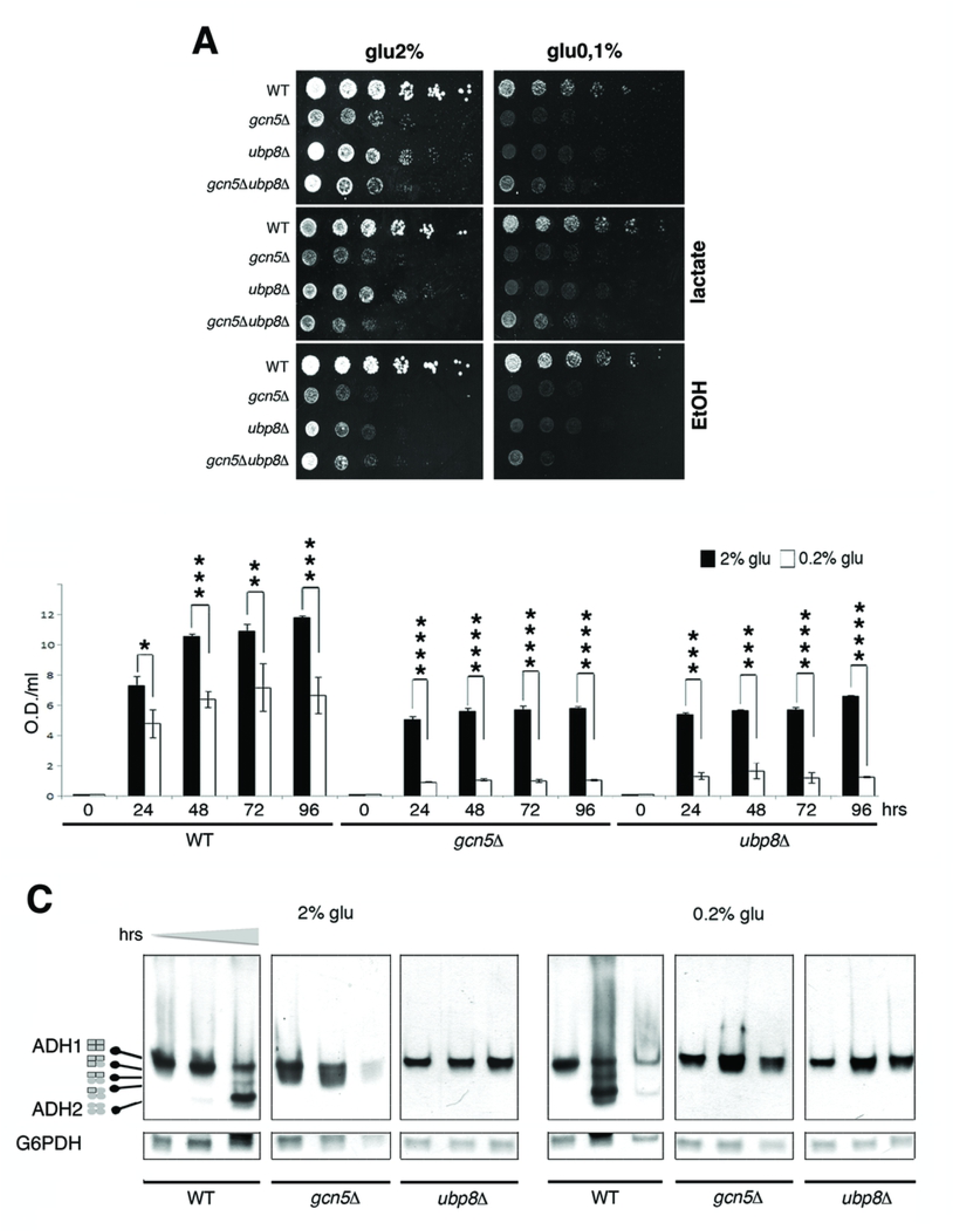
Loss of Gcn5 and Ubp8 causes defects in glycolysis with poor growth in low sugar. A, Growth spot assay on medium containing high (2%) and low (0,1%) glucose shows defects in absence of Gcn5, Ubp8 and both. Lower panels sho also that poor growth can’t be rescued by addition of downstream glycolytic products such as lactate and ethanol indicating the severity of glycolytic defect. B, Comparison of growth in 2% (black) and 0,1% glucose (white) of WT, *ubp8Δ* and *gcn5Δ* strains in liquid cultures at increasing times. C, In-gel native assay and staining of ADH1, ADH2 extracted from wt, *gcn5Δ* and *ubp8Δ* strains grown in 2% and 0.2% glucose for 30, 60 and 90 hours respectively. The same gel was successively stained for G6PDH shown as separate lane. was performed for sample normalization. Outlined scheme on the left describes the differential composition of Adh1 and Adh2 tetramers shown in the gel.

## Discussion

A complex code of post-translational modifications is responsible for the dynamic regulation of chromatin and for the function of non-histone proteins in every cellular compartment. PTMs are able to modulate therefore the activity of enzymes involved in metabolism where, the transcriptional regulation cannot completely explain the ready response of the cell to sudden changes of external stimuli. For this reason, in short term response it was postulated a key role of post-translational regulation. In yeast, the control of sugar metabolism, glycolysis and fermentation shows nodal enzymes marked by phosphorylation, acetylation and ubiquitylation [11] and the acetyltransferase Gcn5 and Ub-protease Ubp8 were shown to play a role in respiration [12] [6]. The present investigation was undertaken to gain insights in the effects of SAGA components on the ubiquitylation of total proteins. We carried out a comparison of the Ub-proteins identified by a proteomic analysis in strains disrupted in Gcn5, Ubp8 or both in comparison with wild type. To this end, we employed the µLC-MS/MS system with which we first characterized H6-Ub-proteins obtained from the four examined strains and then we achieved quantitative information on specific proteins belonging to cellular metabolic processes subjected to fast changes in response to genetic or environmental stimuli. In fact, as stated above, unlike transcriptomics, our proteomic approach was able to extract the trend of relative abundance of differentially expressed proteins among the three mutated strains compared to WT strain. In addition, it was possible to construct a complete proteomic interaction map to better visualize the major pathways in which these proteins resulted involved. In the first instance, the results obtained clearly indicated a central role for Ubp8 and Gcn5, affecting specific proteins that contribute to the control of cell growth, stress response and above all energetic metabolism. The comparison of Ub-proteins in WT, *gcn5*Δ, *ubp8*Δ and *ubp8Δ*/*gcn5Δ* strains identified proteins whose ubiquitylation vary significatively thus indicating their *bona fide* targets. Moreover, most of the Ub-proteins found as differentially expressed in the examined comparisons, although not always with the same levels of abundance, resulted shared and up-represented in absence of Ubp8, Gcn5 and in the double disrupted strain. These evidences can be interpreted as due to modification of lysines that can alternatively be acetylated or ubiquitylated [11], or, even, to a lower expression of Ubp8 in *gcn5Δ* strain [6]. As also reported by Wilson et al. [9], our data confirmed that SAGA modulates the PTMs of SNF1, involved in the inactivation of enzymes of fatty acid biosynthesis and glycogen storage and it is a positive regulator of autophagy [21], accordingly and found a lower ubiquitylation of this protein in the *ubp8Δ* strain. After grouping these Ub-proteins respect to their functional category we observed the presence of major enzymes involved in glycolysis. Noteworthy, the same proteins were described to be ubiquitylated/acetylated [11]. Remarkably, Pfk1 and Pfk2 key nodes for the execution of the glycolytic flux [22] remarkably, there was no upregulation at transcriptional level of these genes confirming that the higher degree of ubiquitylation found is at post-translational level. It can not be ruled out that an additional role of ubiquitylation might be also involved in a previously proposed allosteric regulation of the enzymatic activity of Pfk1 and Pfk2 [23]. Other genes determining the final utilization of glucose, were identified in our screening (Tab1 and Fig. 3). In the presented model we indicate alternative metabolic redox-balancing routes undertaken in absence of Gcn5 and Ubp8. We report a biochemical analysis of strains growth in high and low sugar where Adh1 and Adh2 activities were used as metabolic marker of fermentation and respiration [24]. Growth was severely impaired in 0,2% glucose accordingly, an extremely low fermentative potential was correlated to the exclusive presence of Adh1 (Fig. 5C) in *ubp8Δ* strain demonstrating its full inability to switch to respiratory conditions as reported in a previous reports [6]. On the contrary, *gcn5Δ* strain displayed weak amount of Adh2 in fermentative conditions (2% glucose) suggesting the rerouting of the glycolytic flux towards a respiro-fermentative metabolism (Fig. 6). We believe that this is probably due to a partial reoxidation of Adh1-dependent accumulation of ethanol by small amount of Adh2, sufficient to transfer the cytoplasmic NADH redox excess to the mitochondrial compartment outer mitochondrial membrane transdehydrogenases (Nde1 and Nde2) involved in glycolytic fermentation to ethanol (Fig. 6) [25]. In this respect we suggest that the differential capacity of external and internal NADH dehydrogenases in the two deleted strains, interpreted in terms of redox metabolism, plays crucial roles in triggering of fermentation [20]. In low 0,2% glucose there is a complete lack of Adh2 that is usually regulated by Adr1 repressed in high glucose [26]. In addition to Pfk1 and Pfk2, showing highest levels ubiquitylation in absence of Gcn5 and Ubp8, other important enzymes such as Tdh2, Pyc1 and Cdc19 at the crossroads of glycolysis, gluconeogenesis and pentose phosphate pathways, showed altered ubiquitylation in both deleted strains (Fig. 4A, B). On the basis of our collected results, we can speculate, that an altered ubiquitylation affects the function of many enzymes involved in sugar metabolism leading to the utilization of an altered redox balance response in absence of Gcn5 and Ubp8. A schematic model, presented in Fig. 6, summarizes our collected results and propose alternative metabolic pathways induced by altered ubiquitylation occurring on major enzymes involved in sugar metabolism in absence of Gcn5 and Ubp8. The presented results open a novel field of investigation and demonstrate the important role of SAGA components Gcn5 and Ubp8 in affecting the ubiquitylation of important metabolic enzymes involved in sugar utilization. Our results also indicate a reciprocal function of Gcn5 and Ubp8 in coregulating the ubiquitylation process. This is not surprising since lysine acetylation often counteract further ubiquitylation suggesting that there is a reciprocal interlaced role of these factors [2]. We think that the presented results may open the way for a novel framework of research with interesting implications for the study of cancer cells where increased glucose utilization and altered glycolytic flux occur independently to oxygen availability [27] [28]. In case of pro-inflammatory stimuli, for example, the induction of a metabolic switch leading to upregulation of aerobic glycolysis [29] lead to an enhanced utilization of glucose which sustains the high rate of cell proliferation and invasiveness [30]. The role of Gcn5 and Ubp8 affecting protein ubiquitylation in yeast is a novel finding shedding light on the importance of the epigenetic post-translational regulation not only for the regulation of nucleosomes accessibility but also at metabolic level for sugar utilization. In sum, we show a direct requirement of the SAGA acetyltransferase Gcn5 and Ub-protease Ubp8 in the regulation of this process strengthening previously data indicating the presence of acetylated and ubiquitylated version among glycolytic enzymes [11]. As a possible translational application of our results we recall the role of the “cancer signature gene” Usp22, hortholog of Ubp8, in aggressive tumors such as kidney and glioma [31]. We might therefore envisage novel strategies that may alter energy supply and lower glycolysis by targeting SAGA components as novel approaches to slow proliferation and invasion of cancer cells.

**Fig. 6.**
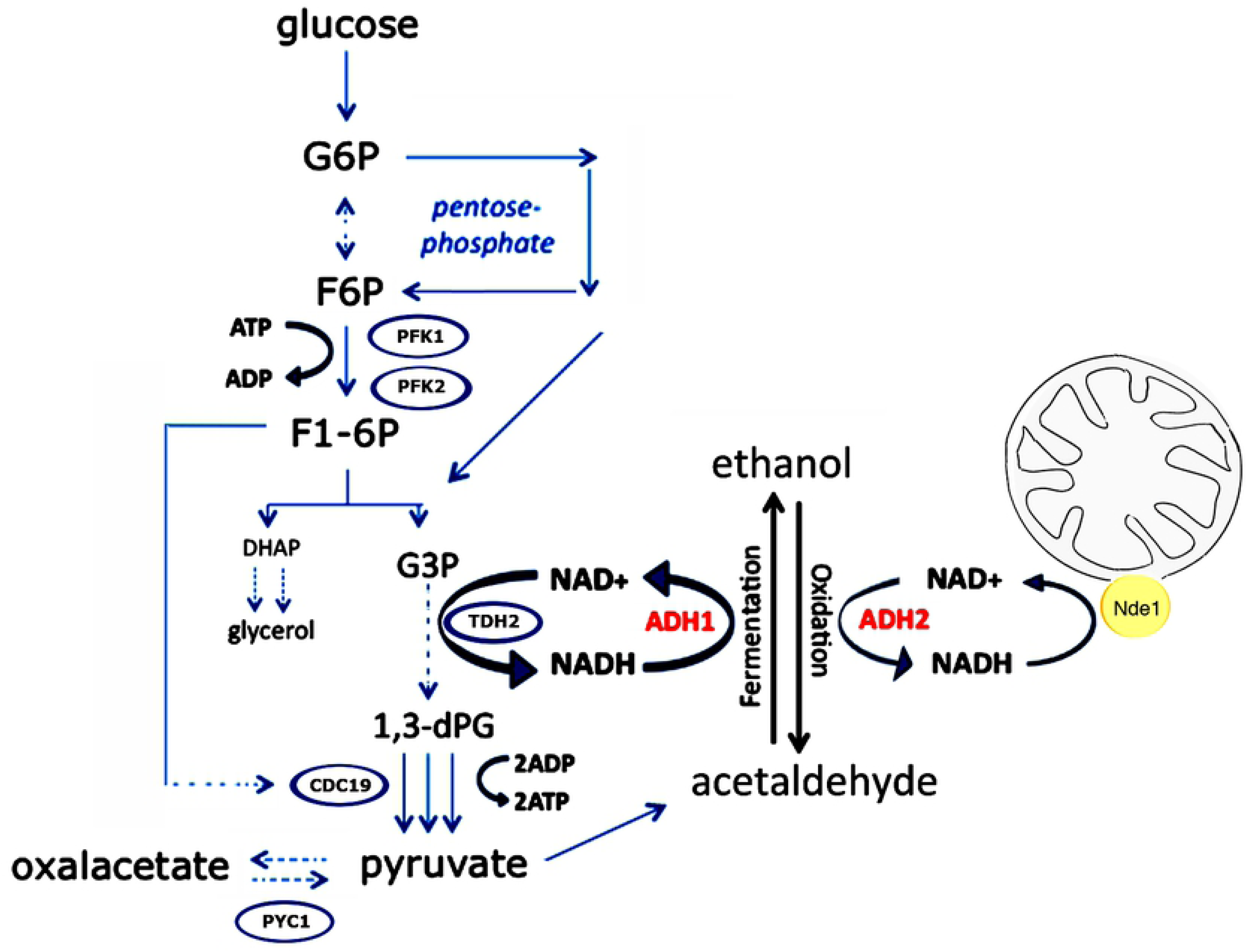
Metabolic effects on glycolysis in absence of Gcn5 and Ubp8. The model outlines enzymes (open blue circles) identified whose ubiquitylation is altered in absence of Gcn5 and Ubp8 and indicates possible routes for the re-equilibrium of NAD/NADH utilization. All the enzymes identified direct the glycolytic flux towards fermentation, gluconeogenesis/anaplerotic routes or pentose phosphate shunt according to the altered redox-balance of the mutants. Cdc19 is regulated by an allosteric modification activated by F1-6P (see arrow).

## Materials and Methods

### Yeast strains and growth

*Saccharomyces cerevisiae* strains derive from the isogenic WT W303 and are listed in Table strains. Gene disruptions of GCN5 and UBP8 or both was carried out with integrative marker cassette and controlled for in situ integration by colony PCR. His6Ub was expressed from pDJ421 plasmid. Growth at 28°C, in YP medium (1% yeast extract, 2% bactopeptone, 1% agar) containing 2%, 0.2%, and 0,1% glucose, 3% glycerol, plus 1% EtOH or 1% lactate. Growth assay: 1 OD_600_ of exponentially growing cells was serially diluted (1/5) and spotted.

### Expression, biochemical purification of his6-Ub proteins

Cells transformed with pDJ421 encoding 6His-ubiquitin under the CUP1 inducible promoter P_CUP1_, [32] were grown on selective media and activated overnight with 0.1 mM CuSO4. Growth was stopped at 0.8–1 OD_600_, cells were lysed in guanidinium buffer (6M guanidinium hydrochloride, 0.1M Na2HPO4/NaH2PO4, 0.01M Tris-HCl, pH8.0, 0.1% Triton X-100, 5mM imidazole, 10mM β-mercaptoethanol, NEM 0.005M, 0.1mM MG132 and protease inhibitors), 1/20th of the lysate is used as input material and TCA (5.5%) precipitated; the rest of lysate was incubated on a Ni-NTA agarose resin (QIAGEN) at 4 °C with rotation. For SDS–PAGE run, blotted and detected of proteins the resin was washed three times with urea buffer (8M urea, 0.1M Na2HPO4/Na2HPO4, 0.01M Tris-HCl pH6.4, 3.5mM β-mercaptoethanol, and 0.1% Triton X-100) before elution in 2X sample buffer (0.125M Tris-HCl, pH6.8, 4% SDS, 0.285M β-mercaptoethanol, 20% SDS, 2% bromophenol blue, 4ml 100% glycerol). For sample preparation for MudPIT analysis, the resin was washed three times with urea buffer 8M and three times with urea buffer 1M and 0.4 μg of trypsin was added to 0.02 μg of the different protein extracts. After an overnight digestion at 37 °C, the reaction was stopped at pH 2.0, by addition of 1 μl of trifluoroacetic acid.

### SDS-PAGE, western blot and immunohistochemistry

Yeast cultures grown at 28 °C semi-complete medium (0,67% yeast nitrogen base, 2% glucose, 0.1% drop out mix). For expression of plasmid 0.1mM of CuSO_4_ was added to culture. Cells were collected at exponential phase (0.8 OD600/ml). Eluates and lysate were loaded on 7.5% SDS–PAGE run and blotted on nitrocellulose membranes (Amersham). Western Blots were hybridized with primary antibodies anti 6His-Ub (Abcam), anti-Ada2 (Santa-Cruz) and anti-AcLys (Santa-Cruz). Proteins were detected by Long Lasting Chemilumiscent Substrate (EuroClone) and visualized by ChemiDoc™ MP Imaging System (Biorad).

### Real time qRT-PCR

Exponentially growing cells (WT, *gcn5Δ* and *ubp8Δ* strains) were collected after overnight growth in YP-glu2%. Total RNA extraction was performed using phenol method and retro-transcribed with QuantiTect Reverse Transcription Kit (Qiagen). Standard curves of WT genomic DNA (20/0,05 ng) and cDNA (50 ng) were amplified for Actin, PFK1, PFK2 and PYC1 with fw and rev primers (listed). All experiments were performed in triplicate. Quantitative RT-PCR (qRT-PCR) experiments were carried out on a Rotor-gene Q (Qiagen) apparatus.

ACT1 Fw: 5’-GCTGAAAGAGAAATTGTCCG-3’

ACT1 Rev: 5’-ACACTTCATGATGGAGTTGTA-3’

PFK1 Fw: 5’-GCTCAAAGTCAAGGTGCTCT-3’

PFK1 Rev.: 5’-CCTGAGAAGGTGATGTTGTTG-3’

PFK2 Fw: 5’-GCAGTTTCAACCAAGCCAAC-3’

PFK2 Rev: 5’-CGTTAGAGTTCATACCTGGAG-3’

PYC1 Fw: 5’-CGTACCGCTCATGAACTGT-3’

PYC1 Rev: 5’-GGATGGTGAAATCTACCTGG-3’

### Protein extraction, estimation, and tryptic digestion for proteomic analysis

As a last step before analysis, the tryptic digest mixtures were desalted using PierceTM C18 spin columns (Thermo Fisher Scientific), according to manufacturer protocol, and resuspended in 0.1% formic acid. LC–MS/MS conditions/Proteomic analysis. Trypsin-digested samples were analyzed by means of a µicro-liquid chromatographic system coupled with a tandem mass spectrometer through a trap-elute configuration. Briefly, 5μl (3µg) of each digested peptide mixture was first loaded, by means of a Micro Autosampler (Thermo Fisher Scientific, San Josè, CA, USA), onto a peptide trap (Zorbax 300 SB-C18, 0.3 i.d. × 5 mm, 5 µm, 300 Å; Agilent Technologies, Santa Clara, CA, USA) for concentration and desalting with a pump running in isocratic mode with 0.1% formic acid in water. Then, the automatic switching of a ten-port valve eluted the trapped mixture on areversed phase column (Biobasic-C18, 0.180 i.d., 100 mm length, 5 μm particle size, Thermo Fisher Scientific) for the separation with an acetonitrile gradient (eluent A, 0.1% formic acid in water; eluent B, 0.1% formic acid in acetonitrile); the gradient profile was 5% eluent B for 5 min, followed by 5–40% eluent B for 45 min, 40–80% eluent B for 10 min, 80– 95% eluent B for 5 min and 95% eluent B for 10 min. The flow rate was 100 µL/min, which was split to achieve a final flux of 2 µL/min. The peptides that were eluted from the C18 column were analyzed directly with a linear ion trap LTQ mass spectrometer equipped with a nano-ESI source (Thermo Fisher Scientific). The spray capillary voltage was set at 1.6 kV and the ion transfer capillary temperature was held at 220°C. Full mass spectra were recorded over a 400-2000 m/z range in positive ion mode, followed by five MS/MS events sequentially generated in a data-dependent manner on the top five most intense ions selected from the full MS spectrum, using a dynamic exclusion for MS/MS analysis. In particular, MS/MS scans were acquired setting a Normalized Collision Energy of 35% on the precursor ion. Mass spectrometer scan functions and high performance liquid chromatography solvent gradients were controlled by the Xcalibur data system version 1.4 (Thermo Fisher Scientific). A total of 16 LC-MS/MS runs were performed, representing the four yeast strain conditions examined (WT, *ubp8*Δ, *gcn5*Δ and *ubp8*Δ/*gcn5*Δ) as technical and biological replicates.

### Mass spectrometry data handling/ Database search

All data generated were searched using the Sequest HT search engine contained in the Thermo Scientific Proteome Discoverer software, version 2.1. The experimental MS/MS spectra were correlated to tryptic peptide sequences by comparison with the theoretical mass spectra obtained by in silico digestion of the Uniprot S.cerevisiae proteome database (6645 entries), downloaded in January 2017 (www.uniprot.org). The following criteria were used for the identification of peptide sequences and related proteins: trypsin as enzyme, three missed cleavages per peptide, mass tolerances of ± 0.8 Da for precursor ions and ± 0.6 Da for fragment ions. Percolator node was used with a target-decoy strategy to give a final false discovery rates (FDR) at Peptide Spectrum Match (PSM) level of 0.01 (strict) based on q-values, considering maximum deltaCN of 0.05 [33]. Only peptides with high confidence, minimum peptide length of six amino acids, and rank 1 were considered. Protein grouping and strict parsimony principle were applied. The output data obtained from SEQUEST software, i.e., Spectral counts (total number of spectra identified for each protein), were treated with MAProMA (Multidimensional Algorithm Protein Map), an in-house algorithm for comparison of protein lists, evaluation of relative abundances, and plotting of virtual 2D maps [14]. The Euler diagrams were calculated using Venny 2.1 drawing tool (software at http://bioinfogp.cnb.csic.es/tools/venny/) were performed by using UniProt accession numbers. Individual cellular function was assigned to each protein according to the GOA database (http://geneontology.org/) and the UniProt database (http://www.uniprot.org/). It should be noted that we were unable to assign the cellular function for some proteins which were therefore classified as “unknown” and additional proteins without a complete characterization at the moment of data analysis were classified as “unclear” or “other”. Label-free differential analysis. The sixteen protein lists obtained from the SEQUEST algorithm were aligned and compared by means of the average spectral counts (aSpCs) corresponding to the average of all the spectra identified for a protein and, consequently, to its relative abundance, in each analyzed condition (WT, *ubp8*Δ, *gcn5*Δ and *ubp8*Δ/*gcn5*Δ). In depth, to select differentially expressed proteins, the four subgroups were pairwise compared, applying a threshold of 0.4 and 6 on the two MAProMa indexes DAve (Differential Average) and DCI (Differential Confidence Index), respectively. DAve, which evaluates changes in protein expression, was defined as (X-Y)/ (X+Y)/0.5, while DCI, that evaluates the confidence of differential expression, was defined as (X+Y) × (X-Y)/2. The X and Y terms represent the aSpCs of a given protein in two compared samples. Most confident up-represented proteins showed a DAve ≥ +0.4 and a DCI ≥ +6; most confident down-represented proteins showed a DAve ≤ −0.4 and DCI ≤ −6. The differentially-expressed proteins resulting from the three most interesting pairwise comparisons (WT *vs ubp8*Δ, WT *vs gcn5*Δ, WT *vs ubp8*Δ/*gcn5*Δ) have been plotted on a protein-protein interaction network built by means of STRING database (https://string-db.org) [15]. Experimentally and computationally predicted interactions were considered for network construction, setting a confidence score of 0.4. Proteins were represented as colored nodes based on their Dave value, highlighting with grey edges their interactions, and were functionally clustered according to their belonging pathway.

### ADH in-gel staining assay

Cell extracts preparation, native polyacrylamide gel electrophoresis (PAGE), ADH and G6PDH staining assays were carried out as previously described [34] [35].

Adh1 and Adh2 protein amounts have been determined from native PAGE with the program ImageJ.

## Aknowledgments

The Authors would like to thank Giuseppe Pisaneschi for technical assistance, PON ELIXIR CNR-BiOMICS (PIR01_00017) to PLM.

## List of abbreviations

WT: wild type
KAT: K-acetyltransferase
DUB: deubiquitylase
PFK1/2: phosphofructokinase
FBA1: Fructose 1,6-bisphosphate aldolase
TDH2,3: Glyceraldehyde-3-phosphate dehydrogenase
CDC19: Pyruvate kinase
PDC1,5,6: pyruvate decarboxylases
ADH1,2: Alcohol DeHydrogenases
ACS2: Acetyl-coA synthetase
PYC1: Pyruvate carboxylase
Dave: Differential Average
DCI: Differential Confidence Index
SILAC: Stable isotope labeling
MAProMA: Multidimensional Algorithm Protein Map

## Supporting Informations Legends

**S1 Table. Complete list of the Ub-proteins detected in WT, *ubp8Δ, gcn5Δ* and *ubp8Δ-gcn5Δ S.cerevisiae* strains.**

In this table for each protein is reported: Uniprot Accession, Reference, Gene name, pI, MW, Spectral Counts (SpCs), Score and Frequency. Frequency indicates how many times a given protein has been identified in the four replicates under each examined condition. The asterisk next to SpC and Score indicates that the average values are given for each protein of the same condition.

**S2 Table. Complete list of differentially expressed Ub-proteins resulted from the MAProMa comparison of *ubp8Δ, gcn5Δ* and *ubp8Δ-gcn5Δ* strains versus WT condition.**

For each protein are reported: Uniprot Accession, Reference, Gene name, Molecular Function, pI, MW, Score, Spectral Counts (SpCs), Frequency, DAve and DCI. Frequency indicates how many times a given protein has been identified under each examined condition. The asterisk next to Score and SpC indicates that the average values are given for each protein of the same condition. Three comparison are considered; WT *vs ubp8Δ*, WT *vs gcn5Δ* and WT *vs ubp8Δ-gcn5Δ.* Positive values for DAve and DCI indicate that the protein is more abundant in the first term of comparison (in this case the WT strain), while negative values for these two indexes indicate that the protein is more abundant in the second term of comparison (in this case the mutant strain considered). For further details regarding the meaning and the confidence range applied to DAve and DCI see Materials & Methods section. It should be noted that proteins were primarly grouped accordingly to their Molecular function and secondarily by their Gene name.

**Table strains.** Complete list of yeast *S.cerevisiae* strains used.

